# Bac-PULCE: Bacterial Strain and AMR Profiling Using Long Reads via CRISPR Enrichment

**DOI:** 10.1101/2020.09.30.320226

**Authors:** Andrea Sajuthi, Julia White, Gayle Ferguson, Nikki E. Freed, Olin K. Silander

## Abstract

Rapid identification of bacterial pathogens and their antimicrobial resistance (AMR) profiles is critical for minimising patient morbidity and mortality. While many sequencing methods allow deep genomic and metagenomic profiling of samples, widespread use (for example atpoint-of-care settings) is impeded because substantial sequencing and computational infrastructure is required for sequencing and analysis. Here we present Bac-PULCE (Bacterial strain and antimicrobial resistance Profiling Using Long reads via CRISPR Enrichment), which combines CRISPR-cas9 based targeted sequence enrichment with long-read sequencing. We show that this method allows simultaneous bacterial strain-level identification and antimicrobial resistance profiling of single isolates or metagenomic samples with minimal sequencing throughput. In contrast to short read sequencing, long read sequencing used in Bac-PULCE enables strain-level resolution even when targeting and sequencing highly conserved genomic regions, such as 16S rRNA. We show that these long reads allow sequencing of additional AMR genes linked to the targeted region. Additionally, long reads can be used to identify which species in a metagenomic sample harbour specific AMR loci. The ability to massively multiplex crRNAs suggests that this method has the potential to substantially increase the speed and specificity of pathogen strain identification and AMR profiling, while ensuring low computational overhead.

**Importance:** There is a critical need for rapid and identification of bacterial strains and antibiotic resistance profiles in clinical settings. However, most current methods require both substantial laboratory infrastructure (e.g. for DNA sequencing), substantial compute infrastructure (e.g. for bioinformatic analyses), or both. Here we present a new method, Bac-PULCE, (Bacterial strain and antimicrobial resistance Profiling Using Long reads via CRISPR Enrichment), which combines CRISPR-cas9 based targeted sequence enrichment with long-read sequencing on the Oxford Nanopore platform. This allows rapid profiling of bacterial strains and antibiotic resistance genes in a sample while requiring very little laboratory or computational infrastructure.

## Introduction

With the rapid increase in antibiotic resistant bacteria, there is a need for methods to quickly identify antimicrobial resistance (AMR) profiles in clinical samples. Previously, most methods of microbial identification and AMR profiling in clinical practice have been culture based (Andrews, 2001; Jorgensen & Ferraro, 2009; Kiehlbauch et al., 2000), an often slow and laborious process.

Over the last decade, a range of newer techniques have been applied for strain and AMR profiling, including whole genome sequencing (Baker et al., 2018), metagenome sequencing (Chiu & Miller, 2019; Gu et al., 2019), mass spectrometry (Havlicek et al., 2013), microarrays (Wilson et al., 2002), microfluidics (Etayash et al., 2016), and others (Syal et al., 2017). A large number of methods rely on the amplification of specific sequences for diagnosis (Jain et al., 2016; Zumla et al., 2014). However, all of these methods require either substantial infrastructure (e.g. for sequencing and analysis); or are effective in identifying a limited range of bacterial strains or AMR profiles.

Recently, Quan et al. 2019 combined CRISPR-based sequence enrichment, PCR, and short-read sequencing to identify bacterial AMR genes in metagenomic samples using a method termed Finding Low Abundance Sequences by Hybridization (FLASH) (Quan et al., 2019). FLASH allows extensive multiplexing for sensitive detection of a wide range of AMR loci while considerably decreasing compute overhead for analysis due to read enrichment. However, there are several limitations of this method. First, the reliance of FLASH on PCR amplification requires enriched loci to be targeted by pairs of crRNAs a specific and relatively short distance apart. Second, the use of short reads makes it difficult to discover linked AMR loci, or AMR context (e.g. plasmid vs. chromosomal). Finally, there are substantial sequencing resources required (although compute resources are considerably reduced).

To circumvent these issues, here we present Bacterial strain and antimicrobial resistance Profiling Using Long reads via CRISPR Enrichment (Bac-PULCE), which combines CRISPR-Cas9-based enrichment of conserved bacterial and AMR loci followed by long-read sequencing (**Fig S1**). We show that this method requires only a single crRNA per target region, greatly increasing flexibility of crRNA design. This method also results in rich information on loci linked to the target sequence of interest, allowing bacterial strain-level resolution even when enriching conserved target sequences (such as 16S). In addition, long reads allow linked AMR genes to be discovered even when they are not targeted.

A critical advantage of Bac-PULCE over long-read metagenomic methods is that enrichment and sequencing of specific sequences from mixed metagenomic samples decreases the computational overhead required for inferring bacterial taxa and AMR profiling. One limiting factor in the use of the Oxford Nanopore sequencing platform is that it requires substantial compute power for basecalling (e.g. GPU) and for downstream bioinformatic tasks (e.g. large numbers of CPUs for read classification). Here we show that it is possible to decrease the amount of sequencing data and computational load of downstream analyses more than 100-fold, while achieving comparable resolution of AMR loci. The efficiency of this method could feasibly allow basecalling and sequence analyses to be performed locally with minimal compute power.

## Results

### Enrichment and sequencing of a variable locus in cultured bacteria

To test the feasibility of using CRISPR-Cas9 sequence enrichment and long read sequencing (Profiling Using Long reads via CRISPR Enrichment; Bac-PULCE) for bacterial strain typing, we first designed two crRNAs (see **Methods**) targeting two sequences surrounding the *E. coli gnd* locus. The *gnd* locus is known to be highly polymorphic in *E. coli*, and has been used previously to type strains (Cookson et al., 2017). We designed one crRNA to target a sequence upstream of *gnd* (within *hisF)* and the other to target downstream of *gnd* (within *wcaM)*, with approximately 20 kilobase pairs (Kbp) between these two target sites (**Table 3**). We then used the Bac-PULCE method to sequence enriched DNA from *E. coli* K12 MG1655 on a single MinION flow cell (see **Methods**).

As a result we generated 43,024 reads, totalling 370.3 Megabase pairs (Mbp) of sequence data with a mean read length of 8,606 bp. The majority of these reads (52.8%) mapped to the *gnd* region. The median coverage depth across the *E. coli* MG1655 chromosome was 33, while the maximum depth at the cut site in *wcaM* was 11,799 (**Fig 1A**). This is an increase of more than 350-fold depth at the target site over background. Importantly, we found that the cutting efficiency of each crRNA differed substantially. The *hisF* crRNA was far less efficient at binding and cutting than the crRNA in *wcaM*. This is clearly evidenced by examining the number of reads that start at each cut site, with greater than four-fold the number of reads originating at the *wcaM* cut site compared to *hisF* (**Fig 1B**).

**Figure 1.**
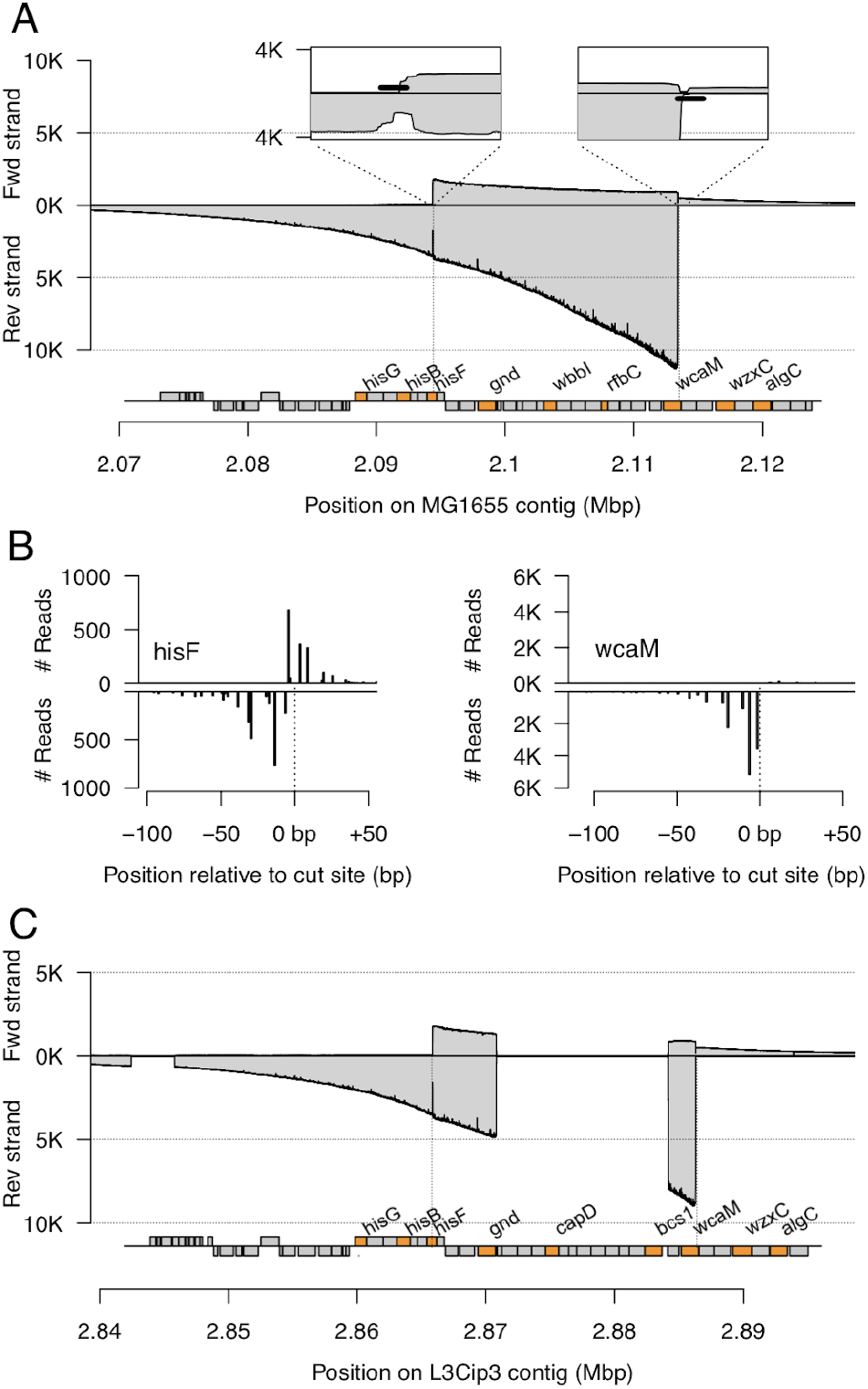
Long reads from Bac-PULCE allow strain-level identification. A. Bac-PULCE allows for more than 100-fold enrichment of sequencing of target loci. We used two crRNAs that flank the highly polymorphic *gnd* locus in *E. coli*, one in *hisF* and the other in *wcaM,* and sequenced the target-enriched DNA using long read nanopore sequencing. We mapped all reads to the *E. coli* MG1655 genome. The coverage depth of reads that map to the top strand is shown above the x-axis, while the coverage depth of reads mapping to the bottom strand is shown below the x-axis. All annotated genes around the *gnd* locus are shown beneath, with several genes labelled for context (labelled genes are coloured in orange). The insets show the cut regions at higher resolution, with the binding sites of the crRNAs indicated by thick lines. **B. crRNAs exhibit clear differences in efficiency and directionality bias**. The two plots indicate the number of reads starting near each crRNA cut site. Lines above the axis indicate reads starting on the top strand; lines below begin on the bottom strand. The left plot indicates the cut site of the *hisF* crRNA. The right plot indicates the cut site of the *wcaM* crRNA. The crRNA targeting *wcaM* is highly efficient and exhibits considerable directionality bias, with almost 10,000 reads originating within 10 bp of the cut site, and these occurring almost solely on the bottom strand. In contrast, the crRNA targeting *hisF* is less efficient, with fewer than 2,000 reads originating within 10 bp of the cut site, and reads starting on both strands. **C. Reads from the *gnd* region of *E. coli* MG1655 mapped to the environmental *E. coli* L3Cip3 exhibit substantial gaps due to loss of homology**. The *wbb* operon region has been replaced in L3Cip3 through a homologous recombination event, resulting in a loss of homology. This suggests that strain-level classification may be possible using long reads from the highly variable *gnd* region.

We also found differences in directionality bias. We expect that the majority of reads starting at a cut site will be in a single direction, despite the fact that the Cas9 cut creates two 5’ phosphorylated ends. This is because the CRISPR-Cas9 complex likely remains bound to the strand containing the target site, and prevents motor ligation and sequencing. However, we found that at the *hisF* cut site, almost equal numbers of reads occurred in both directions (**Fig 1B**). In contrast, at the *wcaM* locus, a majority of reads started in only one direction. Overall, these results indicated that targeting a locus with a single crRNA should allow efficient enrichment through crRNA binding and cutting, followed by long-read sequencing, although crRNA binding and cutting efficiency can differ considerably. We next tested whether reads at the *gnd* locus could be used for strain-level identification.

We first generated genomic sequence data and assembled the genome of a novel environmental isolate of *E. coli* strain, L3Cip3 (Van Hamelsveld et al., 2019), into a single circularised 4.93 Mbp chromosomal contig, four circularised contigs likely to be plasmids (177 Kbp, 88.9 Kbp, 84.0 Kbp, and 44.7 Kbp), and two short circularised contigs likely to be fragments (2255 bp and 1565 bp) (see **Methods**). We then mapped the reads generated from Bac-PULCE from the *E. coli* MG1655 *gnd* locus to this second strain of *E. coli*. We found that although these reads mapped, it was readily apparent that they did not map over their full length, as indicated by sudden drops in the coverage depth (**Fig. 1C**). In this case, the drop in coverage depth was due to the loss (via homologous recombination) of an operon present in *E. coli* MG1655 that contained several genes active in capsule polysaccharide biosynthesis. These results suggested that by using long reads, accurate strain-level classification would be possible using highly variable regions such as the *E. coli gnd* locus, which is prone to homologous recombination. It also suggested that targeting a more conserved gene, such as 16S ribosomal RNA genes would be feasible.

### Enrichment and sequencing of a conserved locus

The *gnd* locus is specific to *E. coli*, and targeting this region in other bacterial taxa, or in a complex metagenomic sample would enrich only for *E. coli* sequences, and thus can not be used to enrich identify sequences from strains of distantly related groups of bacteria. Therefore, we next tested whether accurate strain-level identification would be possible using crRNAs targeting conserved genomic loci. We designed a crRNA targeting the highly conserved 16S ribosomal RNA genes and using Bac-PULCE, enriched and sequenced 16S loci from clonal L3Cip3 genomic DNA.

We generated 78,791 reads, with 92.5% of these reads mapping to the *E. coli* L3Cip3 genome. We also mapped these reads to the *E. coli* K12 MG1655 genome. We found that even when mapping reads that originated from highly conserved 16S loci (of which there are seven total in *E. coli*), in the genomic regions surrounding the 16S loci, small indels and duplications were present that clearly indicated whether reads had mapped as expected (**Fig 2A** and **B**; **Fig S2**). These could only be observed by relying on long reads that extended beyond the conserved 16S locus into these more polymorphic regions.

**Figure 2.**
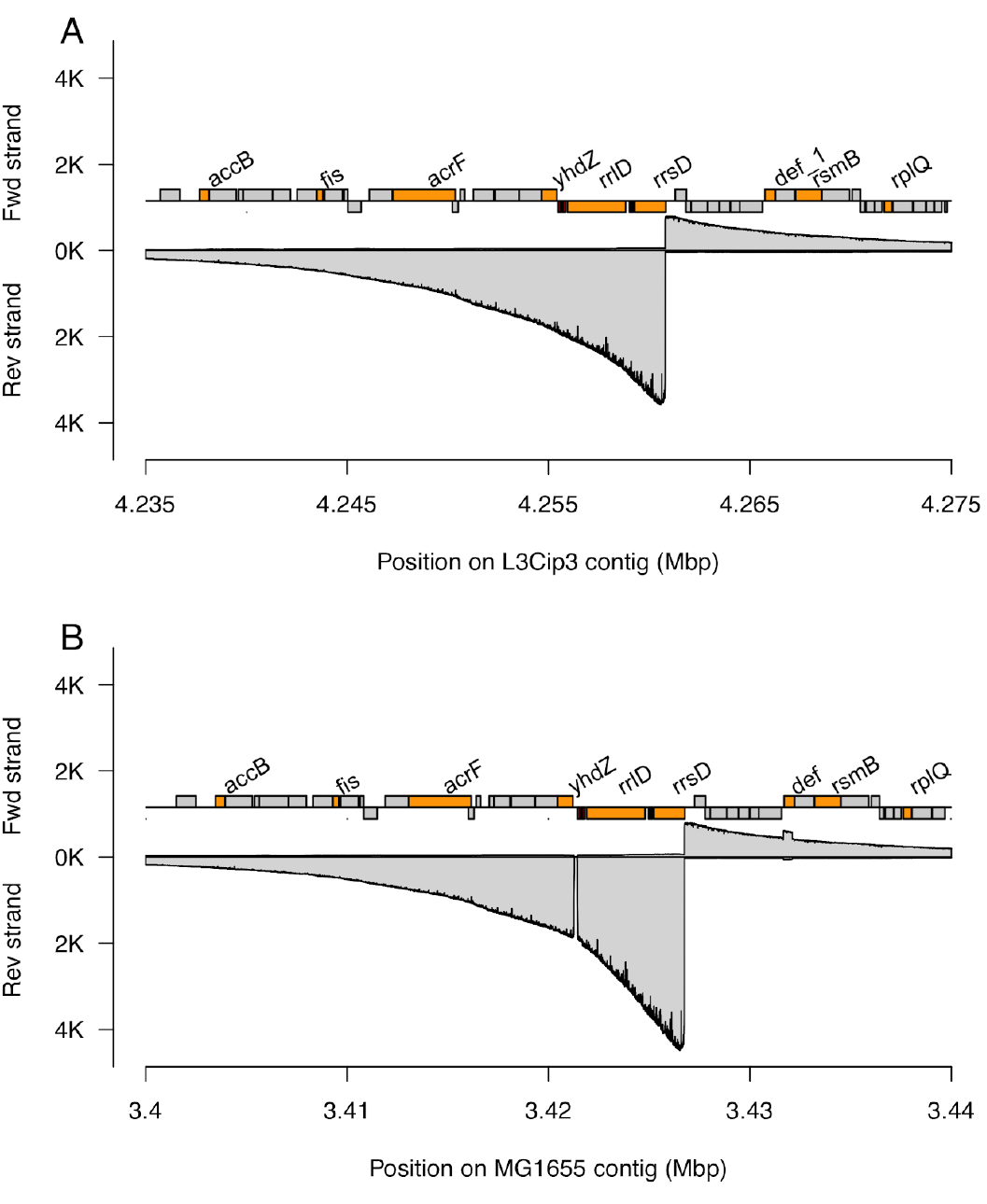
Small insertions and deletions at highly conserved 16S loci allow reads from different strains to be distinguished. A. Coverage depth for one of the seven 16S loci in the L3Cip3 genome. The depth of reads that map to the top strand are shown above the x-axis, while reads mapping to the bottom strand are shown below the x-axis. Several ORFs are annotated (coloured in orange). **B. Coverage depth for the homologous region in the MG1655 genome.** Relative to L3Cip3, there have been small deletions (here in a region just upstream of two tRNAs) and duplications in the MG1655 genome, indicated by drops or increases in coverage. These have occurred adjacent to the highly conserved 16S region, but are only apparent with long reads extending beyond the highly conserved 16S locus. This locus is one of seven 16S loci; all others exhibit similar discrepancies in coverage depth (**Fig S2**).

To test the limits of taxa identification in a more systematic manner, we mapped the reads originating from the 16S loci in L3Cip3 to the rrnDB 16S database (Stoddard et al., 2015), which consists of full length 16S rRNA from 77,530 bacterial species (see **Methods**). We found that 97.6% of these reads had their primary mapping to ribosomal sequences from either *Escherichia* or *Shigella* (which is a polyphyletic genus within the *E. coli* species complex). The vast majority of incorrect matches were short alignments: 99.8% of all mappings with alignments longer than 1400bp were to ribosomal sequences from either *Escherichia* or *Shigella.* These results suggest that using Bac-PULCE to selectively sequence 16S regions allows precise identification of taxa to at least the level of genus.

#### Strain level identification

We next tested the accuracy of using reads from both the *gnd* and 16S loci for strain-level identification. We mapped the *gnd* and 16S reads from *E. coli* L3Cip3 against a database consisting of the L3Cip3 genome and whole genome sequences from 58 additional *E. coli* strains encompassing the diversity of the *E. coli* clade (see **Methods; Fig. S3**). For both *gnd* and 16S, we found that the mapping was highly specific, with approximately 90% of all 16S reads having their primary mapping to the strain of origin. In the case of *gnd*, this fraction exceeded 99% (**Fig. 3**). Furthermore, there was a clear relationship between both mapping quality and read length on the accuracy of strain-level assignment: long reads and reads with high mapping quality were very likely to correctly identify the strain, with accuracy considerably exceeding 99% for reads exceeding 15 Kbp in length even for the 16S locus. This clearly indicates that even when using Bac-PULCE to target highly conserved loci such as 16S rRNA genes, it is possible to precisely identify the bacteria at the strain-level. This vastly improves taxonomic resolution beyond what is currently possible when sequencing just the 16S region, and is made possible by the length of the reads (Johnson et al., 2019).

**Figure 3.**
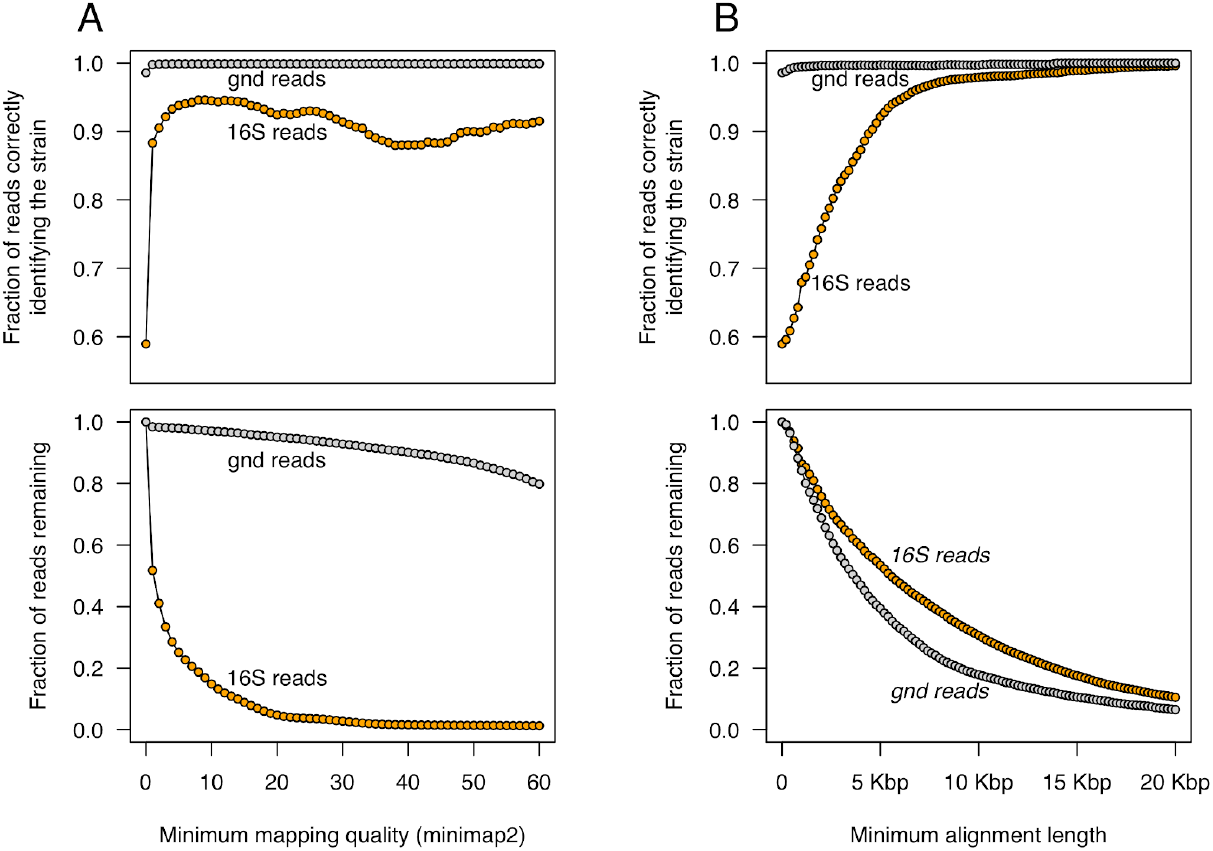
Long reads allow unambiguous identification of *E. coli* taxa at the strain level. A. Relationship between mapping quality and the accuracy of strain assignment for the *gnd* and 16S loci. The top panel shows the fraction of reads mapped to the correct strain (L3Cip3) as a function of mapping quality, while the bottom panel shows the fraction of reads with that mapping quality or higher. At a minimum mapping quality of 1 almost 90% of all 16S reads map to the correct strain despite this locus being highly conserved. The fraction or correctly mapped reads is far higher for the polymorphic *gnd* locus. **B. Relationship between read length and the accuracy of strain assignment.** The top panel shows the fraction of reads mapped to the correct strain (L3Cip3) as a function of read length, while the bottom panel shows the fraction of reads of that read length or longer. In contrast to the relationship between read quality and classification accuracy for 16S, at long read lengths (e.g. more than 15 Kbp), the accuracy of strain assignment exceeds 99%.

#### Sequence context affects Bac-PULCE efficiency

The data here show that accurate strain-level classification is possible even when targeting highly conserved loci. In addition, we found that binding and cutting efficiency can differ substantially between loci. We hypothesised that these differences could arise either from the specific target sequence, or from the genomic context of the target site. We thus next examined variability in the cutting efficiency of different loci for an individual crRNA (Liu et al., 2016).

We designed a crRNA targeting identical sequences in multiple copies of beta-lactamase genes which were present on two plasmids in L3Cip3. All three of these copies are identical in sequence. Here again we found that the crRNA cut with high efficiency and specificity, but that this varied among cut sites both in terms of strand bias and efficiency (**Fig. 4A** and **4B**), despite the target sequences being identical. Of the three beta-lactamase loci targeted by the crRNA, one cut such that Oxford Nanopore motors ligated to both strands at almost equal levels (**Fig 4C**), while a second cut such that reads were phosphorylated almost exclusively at only one end. Again, we hypothesise that directionality bias is due the CRISPR-Cas9 complex remaining bound to the DNA and subsequently blocking ligation of the motor complex. However, in contrast to the results above, when we repeated this analysis for the crRNA targeting the 16S regions, we found that all seven 16S regions were cut and sequenced almost identically (**Fig. S4**).

**Figure 4.**
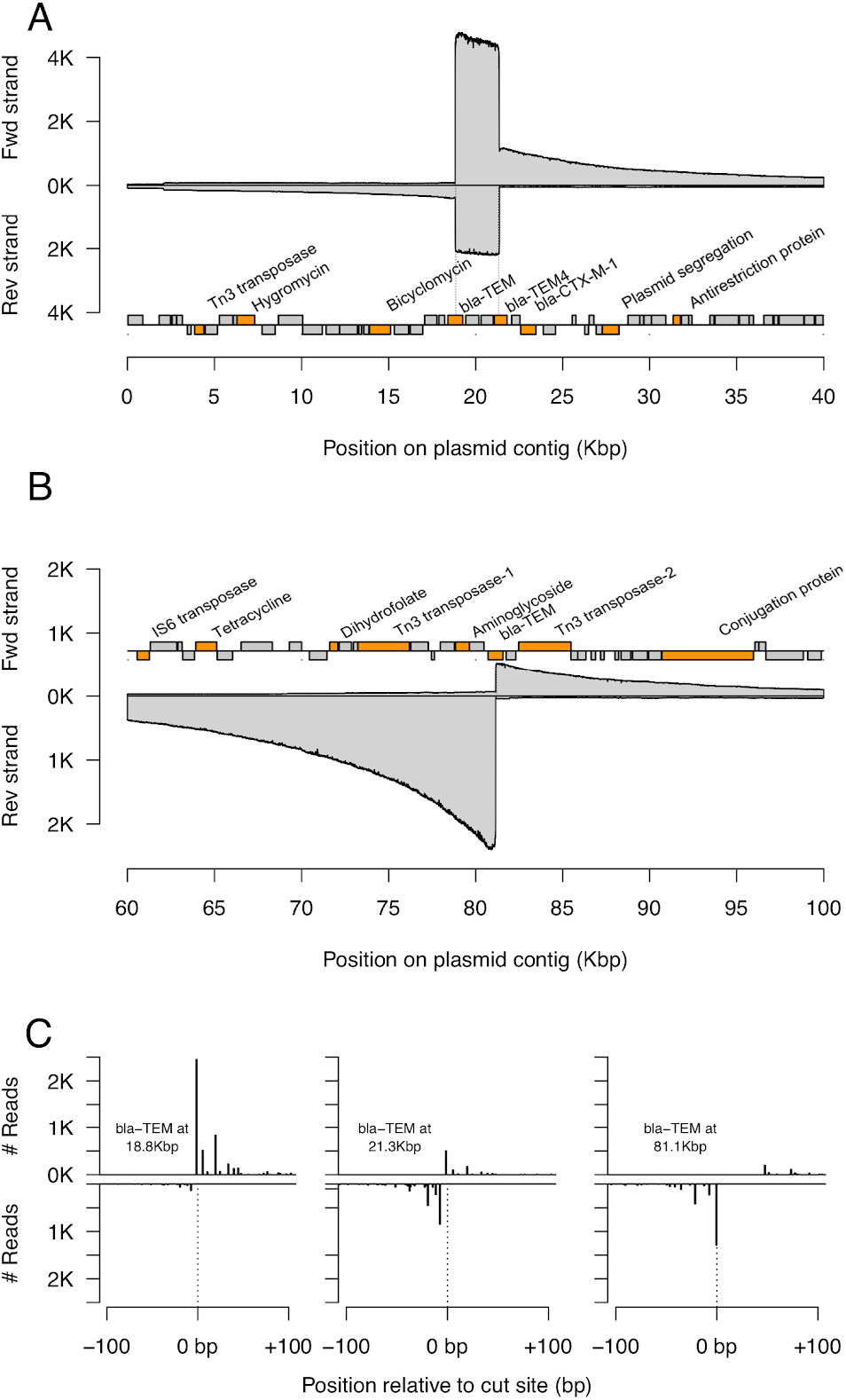
Variability in crRNA cutting for identical target sequences. We used a single crRNA to target multiple versions of a beta-lactamase resistance gene present on two different plasmids in L3Cip3. These target loci have identical sequences, although the surrounding sequence context is different. **A. Coverage depth at a region with two beta-lactamase genes and target cut sites.** The depth of reads that map to the top strand are shown above the x-axis, while reads mapping to the bottom strand are shown below the x-axis, with several ORFs annotated (coloured in orange), including the beta-lactamase resistance genes (bla-TEM). **B. Coverage depth at a region with a single beta-lactamase gene and target cut site. C. Each cut site exhibits a unique binding and cutting efficiency as well as directionality bias**. The three plots indicate the number of reads starting near each crRNA cut site. Lines above the axis indicate reads starting on the top strand; lines below begin on the bottom strand. The plots are shown in the order of cut sites, with the locations of the cut sites indicated on each plot. Cuts at the first bla-TEM locus are efficient and have a clear directionality bias; cuts at the second bla-TEM locus are less efficient and have less bias, with reads almost equally likely to start on the top or bottom strand. Reads at the third bla-TEM locus again show clear bias. This locus is on a separate plasmid that is present at approximately 0.41 lower copies than the first (as inferred through read coverage), suggesting that cutting efficiency differs little between the first and third cut sites.

#### Bac-PULCE allows identification of additional linked AMR loci

In addition to long reads providing information on polymorphic regions near to conserved 16S loci, long reads allowed ready identification of additional AMR loci linked to the targeted bla-TEM loci. One plasmid had only a single copy of the bla-TEM locus, and was thus cut once (**Fig 4B**). However, the majority of the reads extended well beyond this locus, such that an additional three AMR loci were sequenced, including a gene for aminoglycoside resistance, dihydrofolate resistance, and tetracycline resistance. The maximum coverage depth on the bottom strand of the targeted bla-TEM locus on this plasmid at position 81.1Kbp was 2,488. Median read depth was 2,028 at the aminoglycoside resistance locus 1.9 Kbp upstream of the targeted locus; 1,028 at the dihydrofolate resistance locus 9.1 Kbp upstream; and 564 at the tetracycline resistance locus 16.8 Kbp upstream. This contrasts with a median depth of 67 over the whole plasmid. This is also apparent at the level of individual reads. 3,049 reads begin or end within the targeted bla-TEM gene (the majority on the bottom strand). 2,113 (69.3%) of these include part or all of the aminoglycoside locus; 981 (32.2%) contain part or all of the dihydrofolate locus; and 530 (17.4%) contain part or all of the dihydrofolate locus.

### Sensitivity and multiplexing capability

Finally, we tested the sensitivity and multiplexing capability of this method. We first sequenced a metagenomic sample consisting of faecal samples from four sheep and one cow on a single MinION flow cell (see **Methods**), yielding a total of 8.83 million reads and 24.5 Gbp, an amount of data that required more than 24 hours to basecall on standard GPU, and far longer using CPU resources alone. We mapped these reads against the resfinder database (Bortolaia et al., 2020) to identify AMR loci. We found a total of 188 reads matching AMR loci (0.002% of all reads). This varied substantially between samples, from 0.0052% in the single cow sample (124 out of 2.36 million reads) to 0.00071% (5 out of 702 thousand reads) in one sheep sample.

We next designed crRNAs targeting ten different AMR loci found in this metagenomic sample (see **Methods**). Pooling several faecal samples together, we used these ten crRNA and an additional crRNA targeting 16S, and performed Bac-PULCE using a single MinION flow cell. This resulted in a total of 37,200 reads. Of these, 53 reads (0.14%) mapped to four different AMR types (**Table 1**). Some of these were sequenced in numbers close to that of the original metagenomic run (e.g. *cfxA*), despite sequencing approximately 250-fold less data in the Bac-PULCE run. This clearly illustrates the power of this approach, in that far less data is required to achieve a similar level of accuracy in AMR profiling. However, other AMR loci were sequenced far less efficiently or not at all (e.g. the ResFinder loci aac(6’)-aph(2’’)_1_M13771 or aph(2’’)-Ia_2_AP009486, which provide aminoglycoside resistance).

**Table 1.** Number of reads found for different AMR loci. The locus (as annotated in ResFinder) is indicated in the first column, with two exceptions, cfxA and tet(W), for which the specific AMR types cannot be differentiated because the crRNA targets a conserved region in the locus. The number of reads mapping to each locus are indicated in the second and third columns for the full metagenomic run and Bac-PULCE run, respectively, while the last column notes the fold-enrichment of the locus in the Bac-PULCE run.

| AMR locus | Total metagenomic Reads (%) | Total Bac-PULCE reads (%) | Fold-enrichment in Bac-PULCE |
| --- | --- | --- | --- |
| *cfxA* | 80 (0.0009%) | 39 (0.1%) | 116 |
| *lnu(C)_1_AY928180* | 11 (0.0001%) | 11 (0.03%) | 237 |
| *catB_1_M93113* | 7 (0.00008%) | 1 (0.003%) | 34 |
| *tet(W)* | 3 (0.00003%) | 2 (0.005%) | 158 |
| *aac(6')-aph(2'')_1_M13771* | 10 (0.0001%) | 0 | NA |
| *aph(2'')-la_2_AP009486* | 9 (0.0001%) | 0 | NA |
| *nimJ_1_NZ_JH815495* | 2 (0.00003%) | 0 | NA |
| *tet(O)_1_M18896* | 2 (0.00003%) | 0 | NA |

We next aimed to identify the organismal context of these AMR loci, relying on the length of the reads to provide this context. Focusing only on the individual reads that mapped to *cfxA* genes in ResFinder, we used BLAST to find matching taxa in the nt database (**see Methods**), only considering reads with more than 100bp of sequence that was not part of the cfxA gene (36 out of 39 reads). We found that the majority of cfxA genes were contained in a chromosomal context in *Prevotella* spp. (54%; **Table 2**), despite *Prevotella* spp. being present at less than 1% frequency across all samples. Thus, leveraging read length yields considerable insight into the organismal and genomic context of these AMR loci.

**Table 2.** Organismal context of cfxA resistance loci. We trimmed all Bac-PULCE reads mapping to cfxA genes to remove the portion matching the cfxA gene, and BLASTed the read against the NCBI nt database. The genus of the top hit is listed in the first column, followed by the community prevalence of that taxa (inferred through mapping 16S reads to rrnDB), followed by the number of cfxA-linked reads mapping to that genus. Several genera are overrepresented in cfxA-linked reads compared to what would be expected from their community prevalence. The last column indicates the median percent identity for all cfxA-linked reads mapping to that genus. Oxford Nanopore reads have a mean accuracy of approximately 93%, so we would expect a strain-level match to be approximately 93% identical and a species-level match to be slightly lower. Most matches have 90% or less identity; thus we identify taxa only at the level of genus.

| Genus | % of bacterial community | % of cfxA-linked reads (number) | median % identity of cfxA-linked reads |
| --- | --- | --- | --- |
| Prevotella | 0.78% | 54% (21) | 90.0% |
| Porphyromonas | 0.12% | 8% (3) | 81.0% |
| Tannerella | 0.40% | 8% (3) | 89.0% |
| Bacteroides | 6.98% | 5% (2) | 80.9% |
| Capnocytophaga | 0.05% | 5% (2) | 91.3% |
| Lachnospiraceae | 2.88% | 5% (2) | 96.4% |
| Chryseobacterium | 0.02% | 2.6% (1) | 96.4% |

Finally, we quantified the efficiency of 16S enrichment from the metagenomic sample. We first mapped all reads from the full metagenomic run to the rrnDB 16S database. For the full metagenomic run, 61,779 reads (0.70%) mapped to this database. As alignment length is closely related to the accuracy of taxon matches, we filtered this set to consider only read alignments longer than 1200bp (near full length 16S matches (Cuscó et al., 2017). This resulted in 17,257 reads (0.20%). In the same Bac-PULCE run as above, we obtained 1,127 reads (3.0%) matching the 16S rrnDB, with 692 (1.9%) of these reads being longer than 1200 bp. This is only 4% of the total full length alignments we obtained in the metagenomic run, and suggests that although 16S regions were enriched in this dataset, the efficiency was far below the enrichment for AMR loci.

## Discussion

Here we have shown that by targeting and enriching specific loci using CRISPR-cas9 to cut at a single locus, followed by long-read sequencing (Bac-PULCE), we can profile bacterial taxa at strain-level accuracy. We have shown this is possible using highly conserved 16S rRNA loci, allowing for far greater taxonomic resolution than is currently available from even sequencing the full length 16S gene. We have also shown this method is able to target and enrich, by over 100-fold, sequences from AMR loci in a complex metagenomic sample. Additionally, the long reads generated from Bac-PULCE allow sequencing of unknown loci (e.g. additional AMR genes) linked to targeted regions.

We found wide variation in the efficiency with which different targets were bound and cut by the crRNA. This was most clear when using Bac-PULCE for enrichment of AMR loci from the metagenomic sample: we failed to sequence some AMR loci at all, although up to ten reads were sequenced during the full metagenomic run. In addition to this probable crRNA sequence-dependence, we found that the efficiency of target enrichment depends on the larger sequence context of the crRNA binding site: identical sequences in different genomic locations can differ by more than two-fold in efficiency. The variability we observed emphasises the necessity of optimising crRNA pools for efficient binding, cas9 cutting, and sequencing. This is best illustrated by the inefficient enrichment of 16S loci from metagenomic samples that we observed: despite observing more than 300-fold enrichment of 16S loci in single isolates, we observed only 10-fold enrichment from the metagenomic sample. Further work using large scale multiplexing in complex samples should allow the optimization of crRNA target sites to improve the efficiency of the Bac-PULCE approach.

There are three primary advantages of Bac-PULCE over other CRISPR-Cas9 enrichment strategies and short-read sequencing methods such as FLASH (Quan et al., 2019). First, by targeting AMR loci with single crRNAs, long reads enable sequencing of linked AMR loci, increasing the resolution of profiling even when all AMR genes are not targeted for enrichment and sequencing. This is also advantageous for profiling bacterial strains: highly conserved loci, such as 16S, can be targeted such that a broad range of bacteria can be profiled. By matching these sequences against 16S databases (such as rrnDB), major genera can be profiled. Strain-level resolution can then be obtained by taking the subset of reads that match each genus (or species), and mapping these against genomes from a wide range of strains within this genus (as we have done here). This is a powerful approach, and could allow strain-level resolution of pathogens from complex samples even when the genus or family of the pathogen is unknown.

Second, because sequencing reads can be of any length and no PCR step is used, only a single cut site is required, considerably increasing flexibility. Third, very little sequencing throughput is required for successful strain typing and AMR profiling. This is critical because although Oxford Nanopore sequencing requires very little laboratory infrastructure, there are still considerable demands for compute power. For example, basecalling a single run usually requires more than 24 hours on a standard GPU. Downstream bioinformatic analyses require additional compute power. Thus, we expect that the limited sequencing throughput required for successful strain typing and AMR profiling should allow rapid screening of complex samples using low-cost infrastructure and less than 1/100th of the compute resources for both DNA sequencing and downstream analyses.

There are, however, two drawbacks to the Bac-PULCE approach at this point. The first is that it requires substantial biomass. Here we have used samples from pure culture or from fecal samples, yielding μg quantities of DNA. This requirement contrasts with approaches that rely upon enrichment followed by amplification, such as FLASH (Quan et al., 2019). However, we expect that by combining Bac-PULCE with methods of non-specific DNA amplification, such as those used for whole genome amplification, we may be able to considerably decrease the amount of biomass required. Second, sequencing efficiency is low. This is a function of both the rarity of the target sequences in the sample, and the efficiency of crRNA binding, cas9 cutting, and attachment of the motor protein. Again, we expect that we can exploit the flexibility of requiring only a single cut site, and the possibility of using highly multiplexed pools of crRNAs to select the most efficient crRNAs for each target sequence of interest (e.g. 16S rRNA).This should further increase the sequencing efficiency and throughput of this approach.

## Abbreviations

CRISPR: clustered regularly interspaced short palindromic repeats
crRNA: CRISPR RNA
Bac-PULCE: Bacterial strain and antimicrobial resistance Profiling Using Long reads via CRISPR Enrichment
AMR: antimicrobial resistance
FLASH: Finding Low Abundance Sequences by Hybridization
Mbp: megabase pairs
Kbp: kilobase pairs
bla-TEM: TEM beta lactamase
GPU: graphics processing unit

## Methods

### DNA isolation

We isolated bacterial genomic DNA from 2mL of overnight cultures using the Promega Wizard Genomic DNA Purification Kit per manufacturer instructions with the following modifications. Following the protein precipitation step, we performed an additional centrifugation step. Additionally, we washed the DNA pellet twice in 70% ethanol. We rehydrated the DNA in 32uL water overnight for 18 hours.

DNA from cow and sheep faecal samples was extracted using the Qiagen PowerSoilPro kit according to manufacturer instructions.

### Genome sequencing

For Nanopore bacterial genome sequencing of L3Cip3 (Van Hamelsveld et al., 2019), we followed the manufacturer’s protocol for the SQK-RBK004 kit (Version: RBK_9054_v2_revM_14Aug2019). We sequenced the sample on a R9.4 flow cell (MinION software MinKnow 3.6.0) and basecalled using guppy v3.4.4. Illumina sequencing was performed by the Microbial Genome Sequencing (MiGS) Center using 150bp PE reads.

### Genome assembly

We used Unicycler v0.4.5 (Wick et al., 2017) for hybrid genome assembly of L3Cip3, with a total of 221 Mbp of Oxford Nanopore data (mean length 2.3 Kbp) and 150bp PE Illumina data (1.99M reads, 525.6 Mbp). We annotated the assembly using prokka v1.14.6 (Seemann, 2014).

### crRNA design

To enrich for the *gnd* locus we targeted conserved sequences in the *hisF* and *wcaM* open reading frames. To enrich for 16S loci we targeted a sequence in *rrsH*, which is present in all seven *E. coli* ribosomal operons. To enrich for beta-lactamase AMR we designed a crRNA that matched all three bla-TEM loci in L3Cip3. To design crRNAs targeting all other AMRs, we used the sequences of the AMR locus found in the ResFinder 4.0 database (Bortolaia et al., 2020).

To design crRNA targeting *gnd*, beta-lactamase, and 16S, we used CHOPCHOP with the CRISPR/Cas9 setting (Labun et al., 2016, 2019), using the human GRCh38 as background. For all other crRNAs, we used the same settings except with *Bos taurus* as background. We set sgRNA length without PAM as 20, PAM-3’ as NGG, allowed up to 3 mismatches in the protospacer, and used the efficiency score from Doench et. al. 2014 (Doench et al., 2014). We filtered all results to retain sequences with GC content between 40-80%, self-complementarity scores of 0, Mismatch (MM) 1 scores of 0, MM2 scores of 0, and MM3 scores <5.

### crRNA and tracrRNA synthesis

We *in vitro* transcribed crRNA and tracrRNA from DNA oligos using a modified *in vitro* transcription protocol (Quan et al., 2019). Briefly, to all crRNA sequences (**Table 3**) we added the T7 RNA polymerase binding site (5’-TAATACGACTCACTATAG-3’) at the 5’ end. To the 3’ end of the crRNA sequences, we added the tracrRNA binding sequence (5’-**GTTTTA**GA**GCTA** TGCTGTTTTG-3’) to allow base-pairing of the crRNA to the tracrRNA.

**Table 3.**
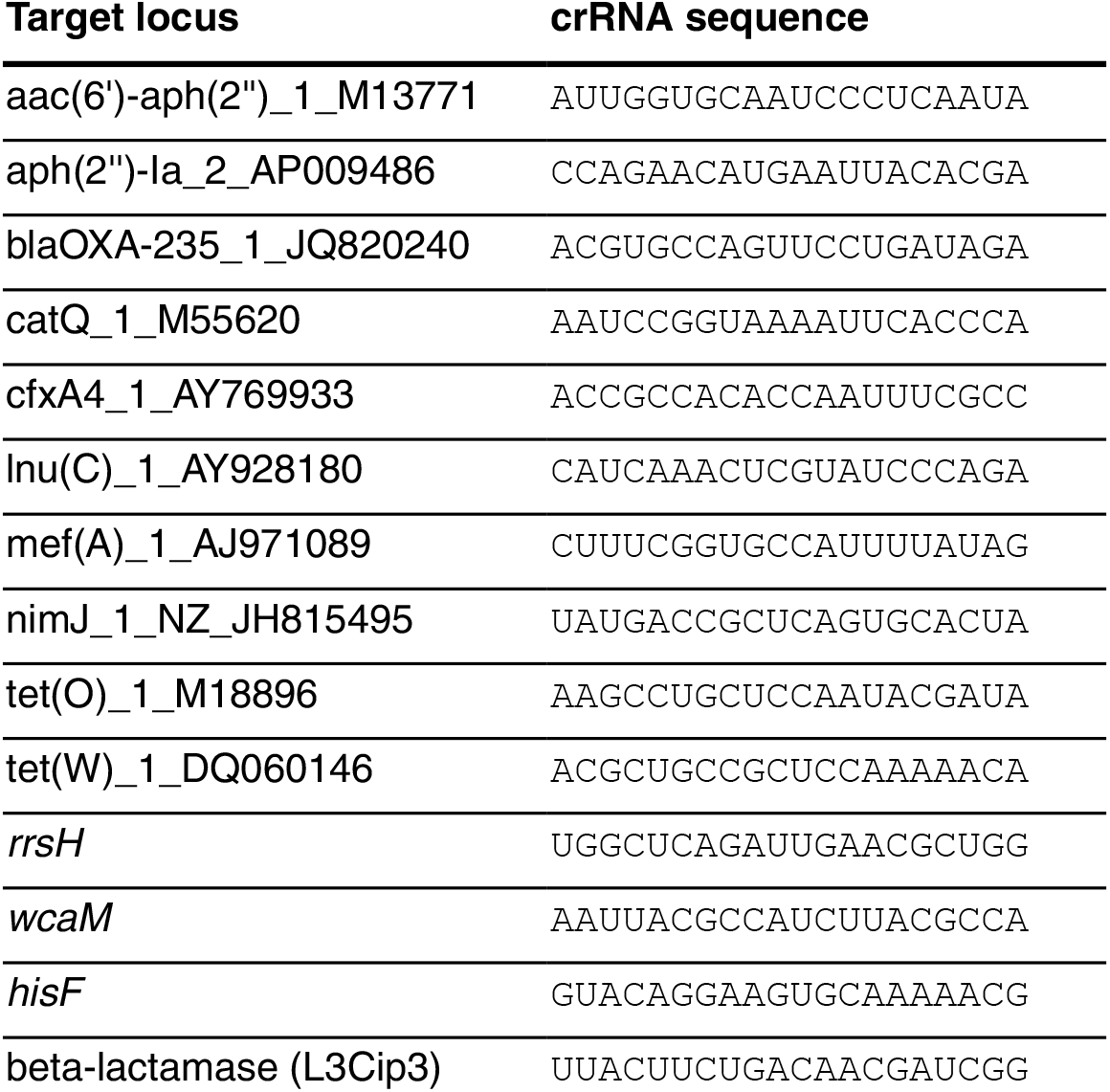
crRNA sequences. The locus (as named in ResFinder for the AMR loci; or the named locus for *E. coli* MG1655) is listed in the first column, and the 5’ to 3’ sequence of the crRNA is listed in the second column. All target regions matching the crRNA have a NGG PAM sequence at the 3’ end.

To transcribe the tracrRNA, we used a DNA oligo with the full length tracrRNA sequence together with T7 RNA polymerase binding site at the 5’ end (underlined). Nucleotides in bold are positions that form base-pairing between the tracrRNA binding sequence and the full length tracrRNA.

5’<u>TAATACGACTCACTATAG</u>GACAGCA**TAGC**AAGT**TAAAAT**AAGGCTAGTCCGTTATCAACTTGAAAAAGTGGCACCGAGTCGGTGCTTTTT 3’

To transcribe the crRNA and tracrRNA from DNA oligos, we used the *in vitro* transcription protocol from Lyden et al. 2019 (Lyden, 2019). up to the step of RNA synthesis. For RNA synthesis we used the NEB Standard RNA Synthesis protocol (E2050, New England Biolabs). We then added 1.5x volumes of ethanol to the reaction, followed by purification using a 1x volume of Ampure XP beads. We eluted the RNA off the beads in 32μL water.

### CRISPR-Cas9 enrichment and sequencing

For target sequence enrichment we used the Oxford Nanopore Cas-mediated PCR-free enrichment protocol v. ENR_9084_v109_revF_04Dec2018 per manufacturer instructions. Briefly, we prepared ribonuclear proteins (RNPs) using pooled crRNAs, tracrRNA, and Integrated DNA Technologies Alt-R S.p. HiFi Cas9 Nuclease V3. We then combined dephosphorylated DNA samples with the RNPs. We dA tailed the CRISPR-Cas9 cleaved target sequences and ligated adapters to these ends.

#### Basecalling and demultiplexing

For basecalling and demultiplexing we used three versions of the Oxford Nanopore guppy basecaller: v.3.2.6 (for the experiment using crRNA targeting *wcaM* and *hisF*); v.3.4.4 (for the experiments targeting *wcaM*, 16S, and beta-lactamase; and for the full metagenomic sequencing); or v.4.0.14 (for the experiment using Bac-PULCE on metagenomic DNA sample). These versions differ by approximately 1% in mean accuracy, and we do not expect that this affects our results here.

### Read mapping and analysis

For all read mapping we used minimap2 with the flags *map-ont* and *--secondary=no*. To test the specificity of mapping for reads originating from the MG1655 *gnd* locus, we considered only reads mapping to a 100 Kbp region surrounding the *gnd* locus in MG1655. To test the specificity of mapping for reads originating from 16S loci, we first extracted reads containing any partial 16S sequence by mapping all reads against all rrnDB sequences from *Escherichia* or *Shigella*. To test for genus-level specificity we then mapped this subset of 16S reads from the sample to the full rrnDB database. To test for strain-level specificity, we mapped the read subsets to a database consisting of 58 whole genomes of *E. coli* (Breckell & Silander, 2020).

To calculate the number of reads originating at the bla-TEM locus that also contained the upstream aminoglycoside, dihydrofolate, or tetracycline AMR loci, we extracted all reads originating within the bla-TEM locus, and mapped these to the open reading frames of the respective AMR gene using minimap2. We inferred that reads successfully mapping to these ORFs contained enough information to determine whether that AMR gene was also present on the read, and thus co-occuring with the targeted AMR locus (in this case, bla-TEM).

To infer bacterial taxa present in the cow and sheep metagenomic samples using 16S reads, we mapped all reads to the 16S rrnDB database. We then filtered all matches to consider only near-full length matches (more than 1200 bp).

To infer the organismal context of the cfxA loci in this complex metagenomic sample, we first identified the reads mapping to any cfxA genes in ResFinder. We then trimmed the portion of the read matching the gene, plus approximately 30 additional bp, and only retained reads with more than 100bp of trimmed sequence. We then BLASTed the remaining portion of each read against a local nt database (downloaded on November 1, 2019).

We performed all statistical analyses using R v 4.0.2 (Stoddard et al., 2015). We performed all visualisations of genomic loci using genoplotR (Guy et al., 2010).

## Availability of data and materials

All read data are available from NCBI (BioProject PRJNA665129). The genome sequence of L3Cip3 is available as BioSample SAMN16242922.

## Authors’ contributions

OKS and NEF conceived and designed the experiments. AS, JW, GF, and NEF performed the experiments. OKS performed the computational analyses. OKS and AS drafted the manuscript, with input from NEF. All authors read and approved the manuscript.

## Acknowledgments

We thank the Heinemann group at the University of Canterbury for providing the L3Cip3 isolate and Dr. Megan Devane of the Environmental Science and Research Crown Research Institute of New Zealand for providing faecal material for metagenomic sequencing.

## Supplementary figures

**Figure S1.**
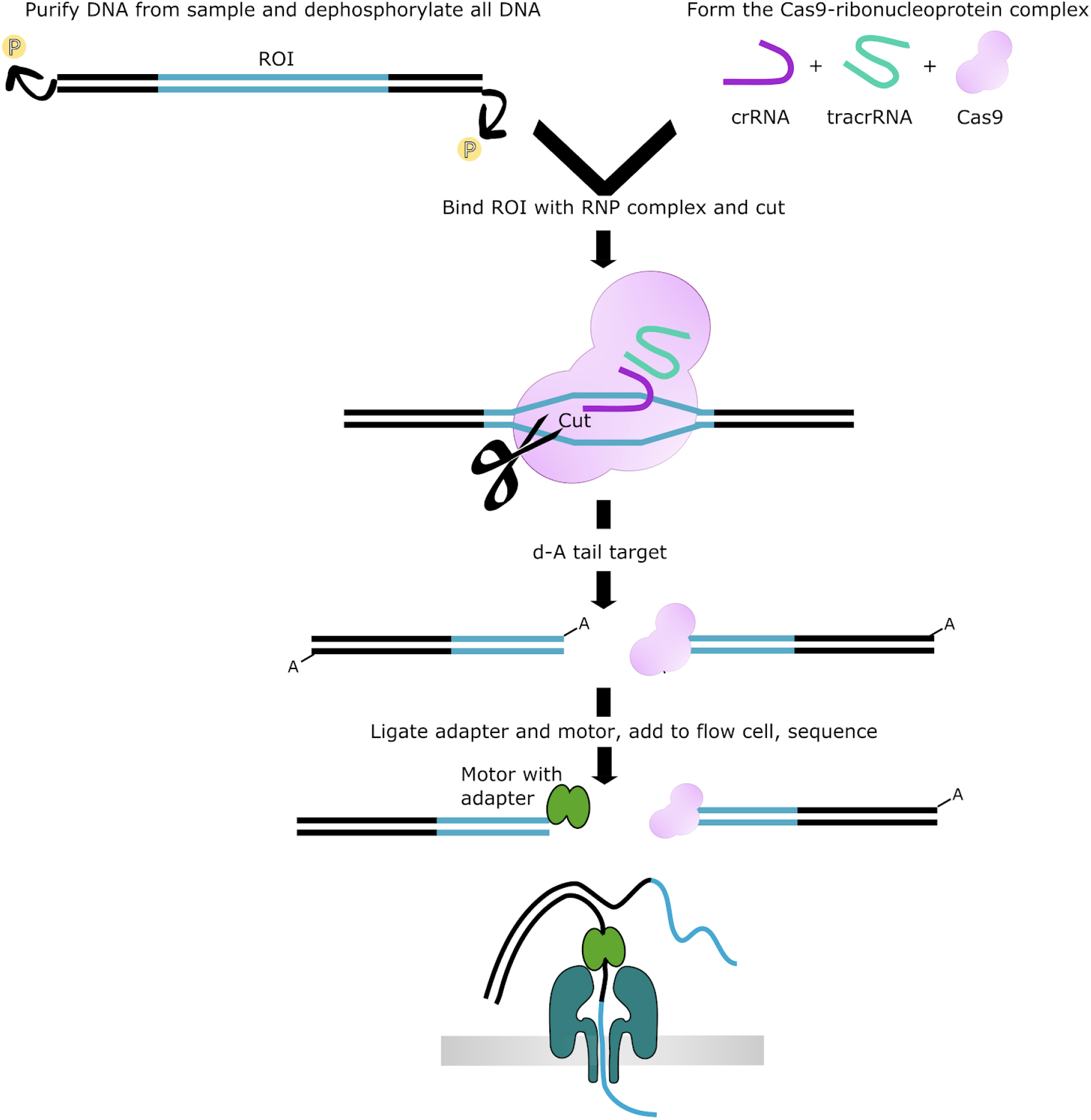
Illustration of the method for CRISPR-Cas9 enrichment and sequencing of targeted regions. First, genomic DNA is dephosphorylated. Ribonucleoprotein complexes (RNP) are formed from cas9, tracrRNA, and crRNA targeting the region of interest. Once cut, the ends of the target are d-A tailed, and sequencing adapters are ligated.

**Figure S2.**
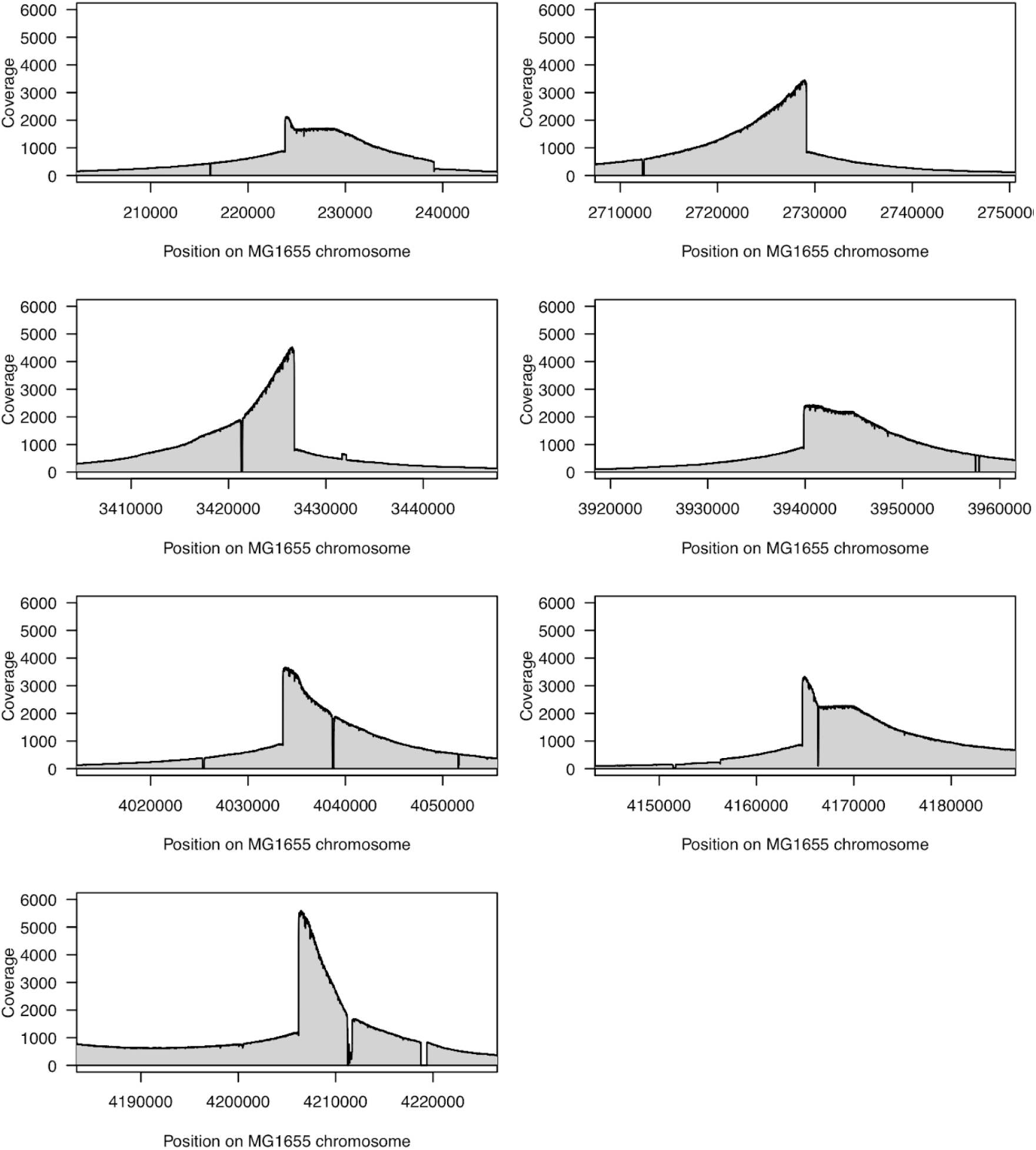
Small insertions and deletions at all 16S loci distinguish *E. coli* L3Cip3 from MG1655. Each panel shows the coverage depth at one of the seven 16S operons in K12 when mapping reads from a Bac-PULCE run targeting L3Cip3 16S with a crRNA. Small insertions and deletions are readily apparent and allow strains to be distinguished when using long reads.

**Figure S3.**
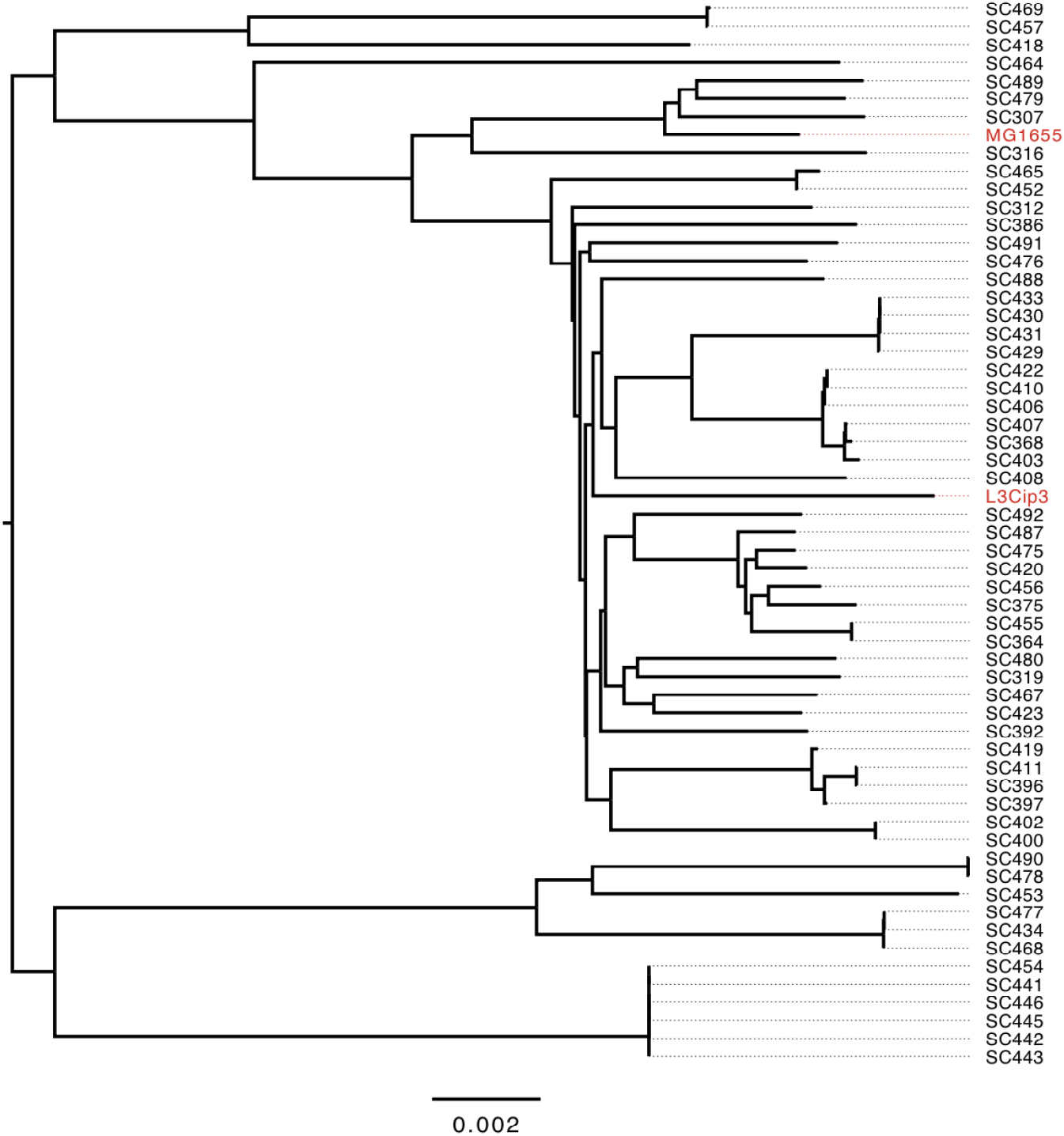
Phylogeny of strains used to test accuracy of strain classification using *gnd* and 16S Bac-PULCE reads. The phylogeny was constructed from whole genome sequences (Breckell & Silander, 2020) using REALPHY (Bertels et al., 2014). *E. coli* K12 MG1655 and L3Cip3 are highlighted in red. This figure was constructed using FigTree.

**Figure S4.**
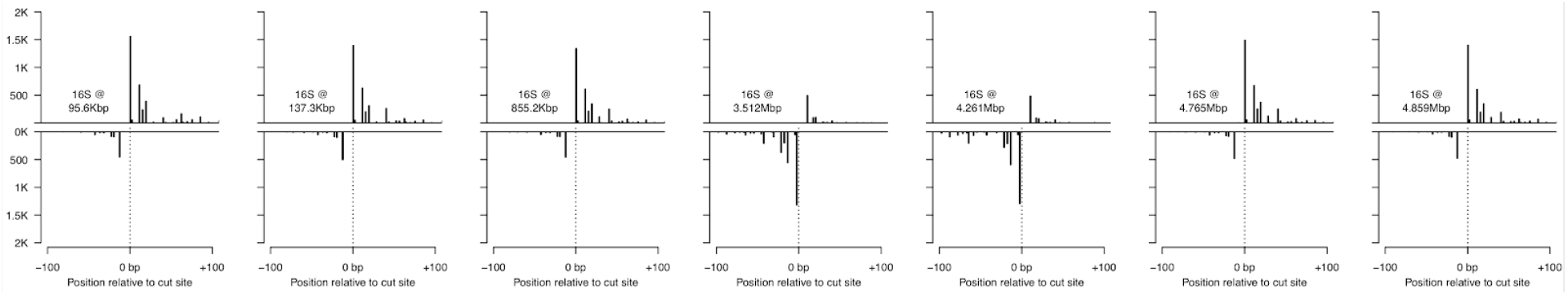
crRNA Binding and cutting efficiency and directionality bias are nearly identical at 16S rRNA regions. The seven plots indicate the number of reads starting near each crRNA cut site. Lines above the axis indicate reads starting on the top strand; lines below begin on the bottom strand. The plots are shown in the order of cut sites, with the locations of the cut sites indicated on each plot. There is very little difference in cutting efficiency or directionality at each site, indicated by the similarity of the sequence start profiles. Reads on the fourth and fifth plots appear primarily on the bottom strand as these operons are organised in the opposite direction compared to the other 16S operons.

